# Origin and dynamics of *Mycobacterium tuberculosis* subpopulations that predictably generate drug tolerance and resistance

**DOI:** 10.1101/2022.10.05.511058

**Authors:** Jees Sebastian, Anooja Thomas, Carly Levine, Riju Shrestha, Shawn Levy, Hassan Safi, Sri Ram Pentakota, Pradeep Kumar, David Alland

## Abstract

Initial responses to tuberculosis treatment are poor predictors of final therapeutic outcomes in drug-susceptible disease suggesting that treatment success depends on features that are hidden within a small minority of the overall infecting *Mycobacterium tuberculosis* (Mtb) population. We developed a multi-transwell robotic system to perform numerous parallel cultures of genetically barcoded Mtb exposed to steady-state concentrations of rifampicin to uncover these difficult to eliminate minority populations. We found that tolerance repeatedly emerged from at least two subpopulations of barcoded cells, one that could not grow on solid agar media and a second that could form colonies, but whose kill curves diverged from the general bacterial population within 4 and 16 days of drug exposure, respectively. These tolerant subpopulations reproducibly passed through a phase characterized by multiple unfixed resistance mutations followed by emergent drug resistance in some cultures. Barcodes associated with drug resistance identified an especially privileged subpopulation that was rarely eliminated despite 20 days of drug treatment even in cultures that did not contain any drug resistant mutants. The association of this evolutionary scenario with a defined subset of barcodes across multiple independent cultures suggested a transiently heritable phenotype, and indeed *glpK* phase variation mutants were associated with up to 16 % of the resistant cultures. Drug tolerance and resistance were eliminated in *ΔruvA* mutant consistent with the importance of bacterial stress responses. This work provides a window into the origin and dynamics of bacterial drug tolerant subpopulations whose elimination may be critical to developing rapid and resistance free cures.

**Importance:** Tuberculosis is unusual among bacterial diseases in that treatments which can rapidly resolve symptoms do not predictably lead to a durable cure unless treatment is continued for months after all clinical and microbiological signs of disease have been eradicated. Using a novel steady-state antibiotic exposure system combined with chromosomal barcoding, we identified small hidden *Mycobacterium tuberculosis* subpopulations that repeatedly enter into a state of drug tolerance with a predisposition to develop fixed drug resistance after first developing a cloud of unfixed resistance mutations. The existence of these difficult to eradicate subpopulations may explain the need for extended treatment regimen for tuberculosis. Their identification provides opportunities to test genetic and therapeutic approaches that may result in shorter and more effective TB treatments.

## Introduction

Unusual among bacterial infections, tuberculosis (TB) requires four to six months of treatment, despite which approximately 5% of cases still relapse (1, 2). Most relapsed TB remains drug susceptible, suggesting that relapse is caused by drug tolerant bacteria which persist during treatment and regrow following treatment withdrawal (3, 4). A peculiarity of TB is that clinical and microbiologic indicators of therapeutic response are poor predictors of a durable cure (5–7). These observations suggest that TB may consist of at least two different *M. tuberculosis* populations: the first, a majority of cells that are responsible for disease symptomatology and culture positivity. This population responds relatively well to drug treatment (3). A second minority subpopulation may consist of cells that are drug tolerant and potentially at risk for becoming drug resistant (8, 9). This subpopulation is responsible for poor TB treatment outcomes, but could be difficult to identify due to its small size or unusual culture characteristics (10). As a transient trait, drug tolerance can be mediated by multiple mechanisms that reduce antibiotic lethality (11–17). Delineating the origin, timing, and unique characteristics of bacterial subpopulations which contribute to drug tolerance in antibiotic exposed *M. tuberculosis* and the pathways by which drug tolerance progresses to drug resistance may inform the development of new anti-tubercular treatments to shorten TB regimen while preventing the emergence of drug resistance. However, the difficulty of isolating drug tolerant subpopulations from either bulk *M. tuberculosis* cultures or clinical TB cases severely limits their study.

Drug tolerance is often studied *in vitro* by exposing bacterial cultures to lethal concentrations of antibiotics and then determining the kinetics of bacterial killing over time (18). However, this approach is prone to artifacts when applied to slow growing bacteria such as *M. tuberculosis* because drug levels can degrade over time, allowing the emergence of preexisting low-level drug resistant mutants in ways that may simulate drug tolerance. Conventional measures of cell survival such as plating for viable colony forming units (CFU) also do not provide insights into the differential survival of clonal subpopulations (10) or the emergence of new subpopulations during drug exposure. We exposed actively growing high-density barcoded *M. tuberculosis* cultures to a steady-state concentration of the anti-tubercular drug rifampicin over a period of up to 30 days, using a Transwell-Tolerance-Resistance (TTR) method to provide us with fine control of drug concentrations in a microplate format. The TTR method is similar to that reported for hollow fiber models (19); however, translating this system into a simplified microtiter plate format made it possible to examine the entire volume of multiple culture wells at each time point and to treat each well as an independent culture that was not conditioned on events that occurred in any other experimental well. These features allowed us to identify different drug tolerant *M. tuberculosis* subpopulations and to trace their propensity to develop drug tolerance as well as the paths taken to eventually emerge with fixed drug resistance.

## Results

### Drug tolerant subpopulations develop before drug exposure

The TTR system’s central components (Fig. 1A) included 24 -well transwell plates, consisting of upper wells for *M. tuberculosis* culture that were each paired with a lower well which acted as an exchange reservoir for fresh media or media-containing antibiotics. Each upper well was separated from its respective lower well by a 0.4 micron membrane permitting the diffusion of the antibiotics while remaining impermeable to cells. Upper wells were loaded with a low inoculum (~ 2 x 10^6^) of *M. tuberculosis*, approximately 2 logs below the number of preexisting rifampicin-resistant mutants measured in mid-log phase seed cultures (8). Steady-state rifampicin concentrations were maintained by robotic exchanges of antibiotic-containing media in the lower wells at defined time points. Equilibration between the upper wells and the lower wells was largely completed after 4 hours (Supplementary Fig. 1A, B) while all bacteria were retained in the upper well (Supplementary Fig. 1C). A specified number of the upper wells (henceforth called wells) were sampled in their entirety at defined time points by direct plating (DP) (on both media alone and media plus rifampicin), and by regrowth plating (RP) as described in Methods. In a preliminary steady-state drug treatment study (Experiment 1) we identified drug concentrations (50X and 20X the rifampicin MIC, corresponding to 0.5 μg/ml and 0.2 μg/ml, respectively) that consistently led to culture sterilization, and found that 10X the rifampicin MIC (0.1 μg/ml) was the highest drug concentration that enabled a subset of cultures to survive up through the final 30-day time point (Supplementary Fig. 2).

**Fig. 1.**
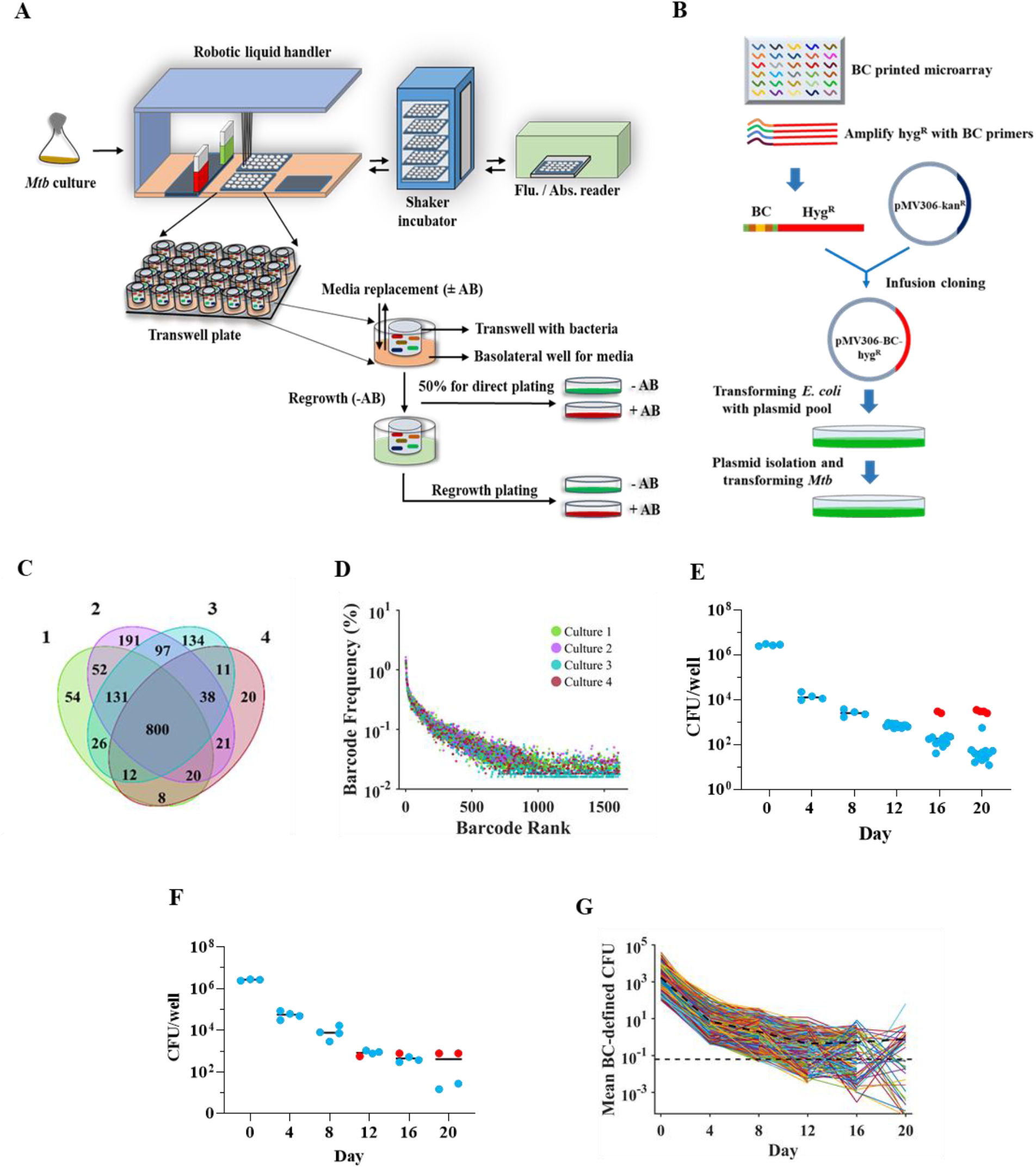
Transwell-Tolerance-Resistance (TTR) system with chromosomal barcoding for high resolution time-kill studies. **A,** TTR system setup. *M. tuberculosis* is seeded into a 24 transwell plate, and a robotic system performs regular media exchanges providing controlled drug levels in each transwell culture. One-half of each well is directly plated onto drug-free and drug-containing media at defined time points while the other half is washed free of drug, regrown to an OD_595nm_ of 0.2 and then similarity plated. **B,** Construction of a barcoded *M. tuberculosis* library starting with approximately 20,000 unique barcode sequences printed on a microarray which produced 4,401 unique barcodes in the subsequently generated plasmid library. **c,** Venn diagram showing the number of different barcodes present in each day 0 culture for Experiment 2 (as indicated by the color legend). **D,** Frequency of each barcode (dots) in each culture (as indicated by the color legend, Y axis). The X axis shows the distribution of all barcodes ranked by frequency. **E, F**, Time-kill kinetic studies using barcoded *M. tuberculosis* cultures showing the number of CFU observed during **E,** Experiment 2 and **F,** Experiment 3 showing rifampicin sensitive (blue dots) and rifampicin resistant (red) CFUs. **G,** Barcode level time-kill kinetics of Experiment 2. Individual barcode count kill curves, adjusted for the number of CFU in each assay well (see methods), denoted as mean barcode (BC) defined CFU, were generated in all replicate culture wells at each time point, excluding wells that contained even one rifampicin resistant CFU. Each line represents the trajectory of a unique barcode. The bold dotted line represents the mean value of total barcodes at each time point. All barcode reads above the 10 barcode count per well cutoff were included in the analysis; however, mean barcodes lower than one is reported (below the dotted line) when this is due to averaging barcode numbers across wells.

We generated a primary stock culture of *M. tuberculosis* that was chromosomally labeled from a library of approximately 4,401 unique barcodes (Fig. 1B) to identify and track bacterial subpopulations across each well and time point. Frozen aliquots of this single primary culture were used to create culture seed stocks for all subsequent experiments to ensure reproducible barcode representation. A seed stock tested at day 0 (prior to rifampicin exposure) of our next TTR experiment (Experiment 2) revealed 1,615 different barcodes across four DP wells, with an average of 1,158 barcodes per well. Barcode composition was relatively well conserved between wells, with 800 of the 1,615 total day 0 barcodes shared by all four day 0 wells (Fig. 1C). Although a few barcodes were more abundant, the great majority of were present at frequencies <0.2% of the total (Fig. 1d).

We used our TTR method to expose and then analyze multiple wells containing culture aliquots of the barcoded library incubated with a non-sterilizing concentration of 10X MIC rifampicin for a period of 20 days. To account for any inter-experimental variability, two separate experiments, Experiment 2 (Fig. 1E) and Experiment 3 (Fig. 1F) were performed, each with identical methods except for differences in the number of test wells at each time point. For both experiments, the DP portion of each well plated demonstrated a two phase kill curve with an initial rapid killing phase characterized by an approximate 0.5 log to 1.0 log drop in CFU through approximately day 12, transitioning to a second persistence phase with delayed killing during days 12-20. This second phase was accompanied by the development of rifampicin resistance by day 16 in Experiment 2 and day 12 in Experiment 3 in a minority of experimental wells; however, the majority of wells from the persistent phase did not produce any CFU when plated on rifampicin (Fig 1E, F), suggesting that the persistence phase was caused by the development or unmasking of drug tolerant subpopulations as has been shown to occur during drug treatment (20). We questioned whether such tolerance was induced at equal rates across all clonal variants within a log-phase culture or instead developed in only a few clonal populations. We used deep sequencing to examine the individual “kill curve” of each clone or small subset of clones marked by a unique barcode in Experiment 2. Normalizing each barcode read count for the number of CFU (mean barcode defined CFU, see methods section) detected in each sequenced well generated high resolution two phase kill curves at the level of the barcode-defined clonal group. In the early stages of rifampicin treatment, the individual kill curves of each barcode defined CFU mirrored the kill curve of each culture as measured by CFU plating in both experiments 2 and 3. Fig. 1G and Supplementary Fig. 3 show that between day 0 and day 12 of rifampicin treatment the number of CFU represented by virtually every unique barcode fell at approximately the same rate as the overall CFU count in each well. However, the trends of the mean barcode defined CFU count and the CFU count per well diverged substantially after day 12, with some barcode defined CFUs showing almost no decline while the overall CFU count continued to drop, albeit more slowly than during the earlier time points. This is explained by the observation that although 84% and 90% of the barcode defined CFU kill curves present on day 12 barcodes disappeared by day 16 and 20, respectively, a subset of barcodes predominated in the remaining CFU in each well, stabilizing the mean barcode defined CFU between these time points. Thus, barcoding revealed that the flattening of the two phase kill curve as measured by CFU plating actually reflects a complex population dynamics among different subpopulations characterized by the emergence of a small number of rifampicin tolerant clonal subpopulations.

We questioned whether tolerance emerges prior to rifampicin treatment or whether tolerance instead develops *de novo* in each well under the stress of drug exposure. In Experiment 2, antibiotic treatment resulted in a dramatic reduction in the number of different unique barcodes between day zero and day 16, reducing these barcodes from a total of 1615 (across all wells) to just 43 different unique barcodes by day 16. However, the number of unique barcodes detected across all wells decreased much more slowly between day 16 and day 20, and some previously undetected barcodes appeared for the first time on day 20 (perhaps because decreasing barcode complexity by day 20 enabled us to detect particularly rare barcodes in the cultures) resulting in the detection of 28 unique barcodes on day 20. A random elimination of bacteria through this severe bottleneck would be expected to generate an increasingly unique distribution of barcodes in each well as CFU numbers progressively decreased over time. Indeed, a clear trend showing fewer barcodes being shared by multiple wells over time was evident during the day 0 to day 12 killing phase in Experiment 2 (Fig. 2A) and the day 0 to day 8 killing phase in experiment 3 (Fig. 2B). However, the effect of this bottleneck was reversed during the persistent phases of both experiments. Thus, approximately equal or greater numbers of barcodes were detected in >76% of the wells compared to 51%-76% of the wells by day 16 in Experiment 2 (Fig. 2A). This trend continued even more markedly by day 20 in Experiment 2 and by days 12 – 20 in Experiment 3 where approximately equal or greater numbers of barcodes were detected in >76% of the wells compared to both 26%-50% or 51%-76% of the wells by day 20 in Experiment 2 (Fig. 2A) and day 12 – 20 in Experiment 3 (Fig. 2B). The barcodes found in multiple wells on day 20 were also more likely to be found in multiple wells on day 16 compared to any time points during the killing phase (Fig. 2C, Supplementary Fig. 4A-F), again demonstrating a tendency for certain barcodes to survive into the persistence phase as independent events in multiple wells. One possible explanation for the survival of the same barcodes across multiple culture wells by the late time points could be that these same barcodes were highly represented prior to drug treatment. However, the relationship between barcode frequency on both day 20 (Fig. 2D) and day 16 (Supplementary Fig. 4G-K) correlated very poorly with the barcode frequency on day 0, while the barcode frequency of day 4 correlated strongly with that on day 0 (Fig. 2E), similar to other time points (Supplementary Fig. 4L-P). Comparable results were observed in Experiment 3 (Supplementary Fig. 5). Thus, the detection of the same barcodes in multiple wells within and across the day 16 – day 20 time points of the same experiment strongly suggests that barcode representation at these late timepoints is not stochastic. Instead, the results suggest that barcoded clones surviving in day 16 – 20 wells underwent purifying selection favoring subpopulations that had developed heritable drug tolerance or a heritable pre-disposition to develop drug tolerance in either the stock culture that was used to create each seed culture or the seed cultures used to prepare *M. tuberculosis* for aliquoting into TTR transwells prior to any drug exposure. Finally, we compared the barcodes present in the day 20 wells of Experiment 2 versus Experiment 3. The seed cultures for each experiment were started with different frozen aliquots of the same stock culture of barcoded cells. Interestingly, even though the majority of barcodes (74.5% ± 19.5 %) were detected in at least one well of both studies on day 0 (Fig. 2F), a significantly smaller proportion of tolerant barcodes (9.23% ± 6.07%) were detected in at least one well of both studies (Fig. 2G) (*P* < 0.0001, one sample test for binomial proportion). These results suggest that tolerance did not pre-exist in the cells making up the original stock. Instead, tolerance appears to have developed separately during the log phase expansion of each individual seed culture prior to the seeding of individual wells and prior to any drug exposure.

**Fig. 2.**
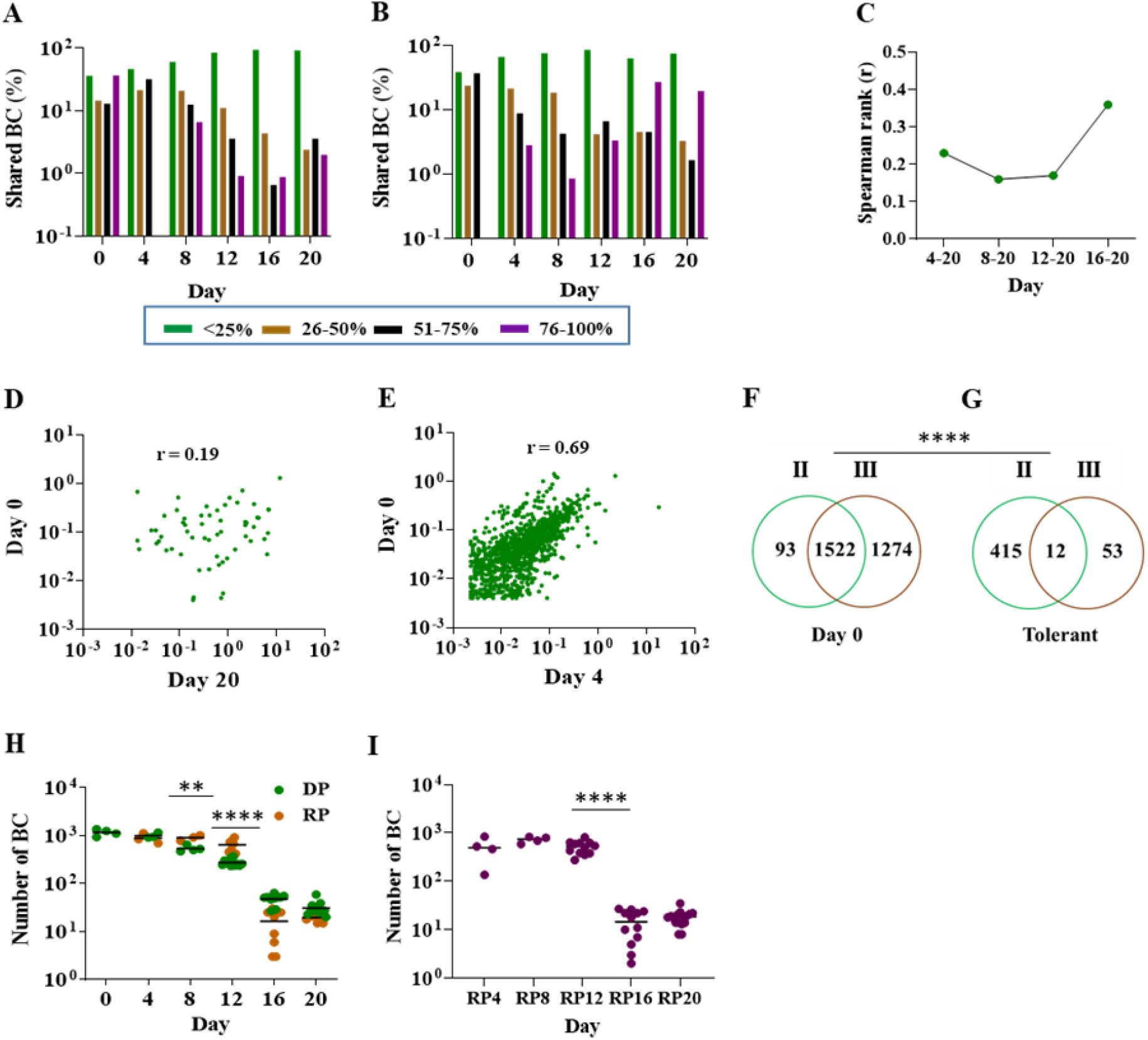
Features of directly plated (DP) and differentially detectable (DD) drug tolerance. **A, B,** The number of barcodes recovered from DP wells that were present in <25%, 26-50%, 51%-75% or >75% of the wells at the indicated timepoint (see key in the figure) for A, Experiment 2 and **B,** Experiment 3. **C,** Trend of the Spearman rank correlations between the number of wells that a barcode is found at day 20 versus each earlier time point, calculated for all barcodes detectible at each indicated timepoint interval. **D,** Correlation of barcode frequency between day 20 and day 0, and **E,** between day 4 and day 0. Each unique barcode that was present in at least one well at each time point is shown by a dot and the frequency for each barcode represents a mean frequency of all wells at that time point. R is calculated by Spearman’s rank correlation. **F, G,** Venn diagram comparing the number of different barcodes in Experiment 2 versus Experiment 3 that were present either in any well at day 0 **F,** or in any well during the drug tolerant phase (days 16 and 20) G (*P* < 0.0001, one sample test for binomial proportion). **H,** The number of unique barcodes identified in Experiment 2 in each well (dots) and at each timepoint. Green dots show barcode numbers of direct plated (DP) cultures and orange dots show barcode numbers after regrowth plating (RP). **I,** The number of unique differentially detectable (DD) barcodes (defined as barcodes that were only present in RP, but not DP wells) identified in Experiment 2 in each well over time. Each indicated time is a regrowth plating (RP) timepoint, which corresponds to the experiment day on which rifampicin was washed from culture wells and allowed to be regrown prior to plating. \**P* <0.05; \*\**P* <0.01; \*\*\*\**P* <0.0001 using a two-tailed paired *t*-test).

### A differentially detectable *M. tuberculosis* subpopulation exhibits transient drug tolerance early after drug exposure

*M. tuberculosis* cultures have been reported to contain drug tolerant “differentially detectable” (DD) subpopulations that cannot be cultured on solid media after drug exposure unless they are first allowed to recover in drug-free liquid media (10, 21). We used regrowth plating (RP) to enrich cultures for DD cells by washing 50% of each well that remained after DP and then re-incubating the wells in fresh media until turbidity indicated substantial growth. The required re-growth period made it impossible for us to determine the original number of DD cells that had been present when the cultures were first washed free of rifampicin. However, barcode counts are likely to be a reasonable although imperfect proxy of this original cell number (Fig. 1G).

After excluding wells that produced even one drug resistant colony on DP or RP to avoid confounding by drug resistance, we found that the number of different DP clones, identified by their individual barcodes, fell at a steady rate up to day 12 (losing ~50% of the number of barcodes at each time point). In contrast, the number of different barcodes detected in RP cultures declined more slowly during this period (Fig 2H), suggesting an increased occurrence of rifampicin tolerance in DD clones at this early drug treatment stage. We then noted a dramatic drop in the number of barcodes detected in both DP and RP samples between days 12 and 16, with very few barcodes remaining in either group of cells by day 16 (43 DP; 17 RP), and a stabilization of barcode numbers between days 16 and 20 (28 DP; 20 RP). To more precisely examine the barcode number trajectory of DD cells, we reanalyzed the regrowth data after removing any of the barcodes detected in DP cultures from the RP barcode count. These data (Fig. 2I) revealed a remarkable stability in DD barcode numbers between days 4-12 and days 16-20 with an approximate 33-fold drop between these two periods. Our results suggest that DD cells exhibit an early but transient form of drug tolerance resulting in their becoming the majority subpopulation early after drug treatment, followed by their return to a minority of cells during the day 16 – 20 drug persistence phase.

### The emergence of drug resistance may be more common in RP than DD cultures

We did not detect any rifampin resistant CFU in any day 0 test well, strongly suggesting that rifampin resistant mutants were absent from all culture wells prior to rifampicin exposure. However, rifampicin resistant mutants did emerge in a subset of DP wells during the persistence phase and in a subset of RP wells starting at day 4 (Fig. 1E, 1F) (Fig. 3A). This finding suggests that the majority of rifampicin resistance emerges *de novo* during drug treatment rather than by the selection of pre-existing drug resistant mutants. Compared to DP wells, RP cultures contained significantly larger numbers of rifampicin resistant clones as indicated by a higher total barcode count detected in resistant cells (93 barcodes versus 28 barcodes for RP versus DP cultures, respectively) and by a larger number of individual wells showing rifampicin resistance after RP culture (18/48, 37%) compared to DP culture (6/48, 12.5%) (*P* = 0.008, Fisher’s exact test) (Fig. 3B). These results suggest that resistance emerges even more frequently in DD than the DP cells; however, the higher number of CFU plated from the RP cultures makes direct comparisons difficult.

**Fig. 3.**
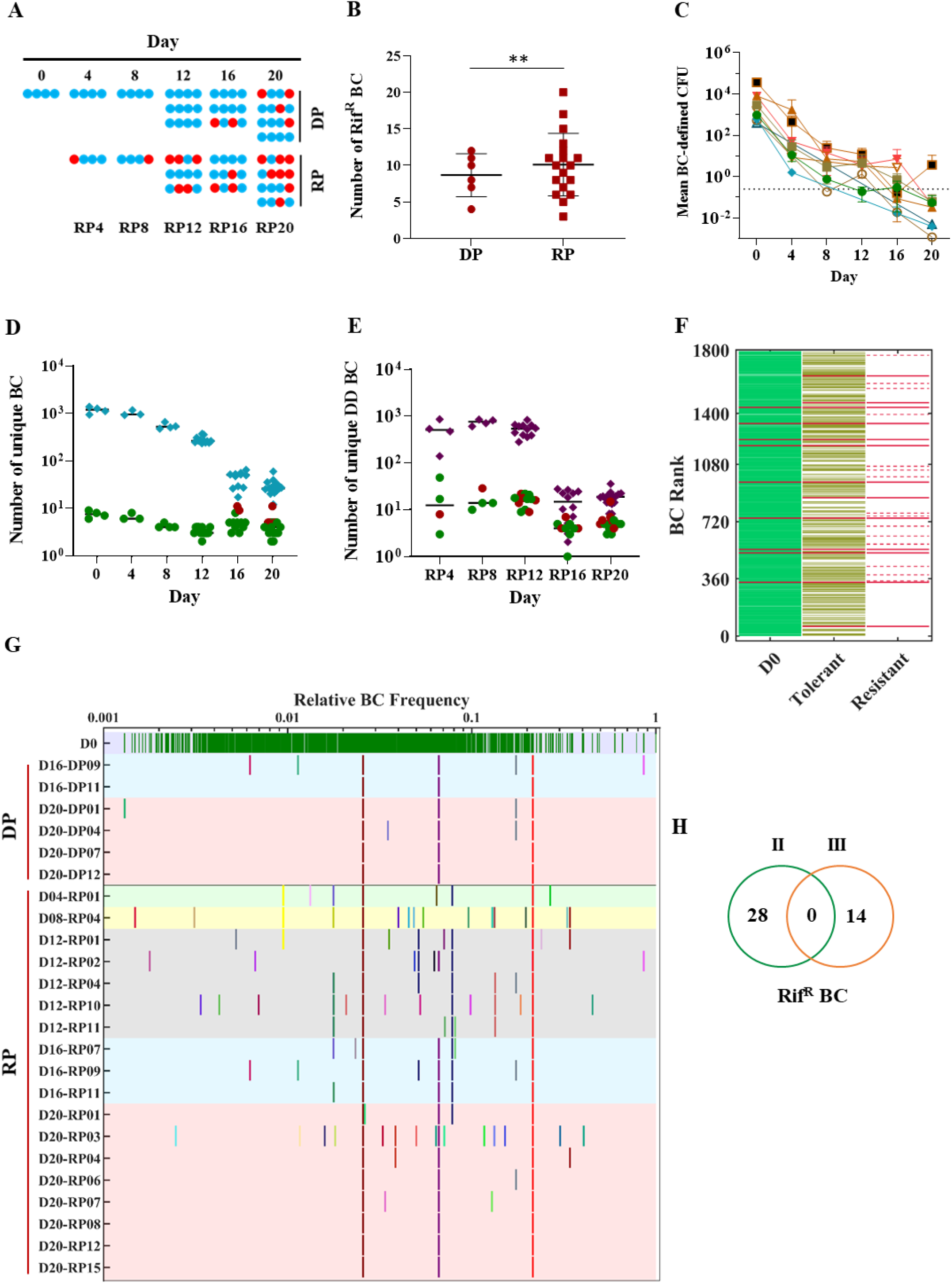
Drug tolerant populations that generate rifampicin-resistant mutants. **A**, Dot plot showing resistance versus susceptible culture results for each well in Experiment 2 plated at the indicated time points. Wells containing only drug susceptible CFU are shown in blue, and those with any drug resistant CFU are shown in red. DP indicates direct plated wells and RP indicates regrowth plated wells with the day indicated corresponding to the experiment day on which rifampicin was washed from culture wells and allowed to be regrown prior to plating. **B**, The number of unique barcodes identified in each well cultured on rifampicin following DP (red circles) versus RP (red squares) (*P* = 0.008, Fishers exact test). **C**, Individual kill curves were generated using deep sequence data of the 9 rifampicin-resistance-associated barcodes identified by DP. All barcode reads above the 10 barcode count per well cuttoff were included in the analysis; however, mean barcodes lower than one are reported (below the dotted line) when this is due to averaging barcode numbers across wells. **D**, The number of unique drug resistance-associated barcodes in each DP well where no drug resistance was detected (green circle) and where resistance was detected by rifampicin plating (red circles) compared to the number of barcodes from these same wells that were not resistance-associated (blue diamonds). **E**, The number of unique drug resistance-associated barcodes in each DD well where no drug resistance was detected (green circle) and where resistance was detected by rifampicin plating (red circles) compared to the number of barcodes from these same wells that were not resistance-associated (purple diamonds). **F**, Barcode plot showing the relationship between each unique barcode detected at day 0 in DP cultures, the barcodes that were still detectable by DP during the day 16 and 20 drug tolerant phase, and the barcodes that were detected by DP in rifampin resistant CFU. Lines on the same vertical axis represent the same barcode. Solid red lines indicated barcodes found in drug resistant CFU which were also detectable as resistance-associated barcodes in drug tolerant or day 0 barcodes. Dotted red lines indicate barcodes in drug resistant CFU that were not detected in either drug tolerant or day 0 barcodes. The order in which barcodes are positioned across the X axis on day 0 is arbitrary. **G**, Diversity and conservation of barcodes detected in drug resistant CFU. All barcodes detected on day 0 are distributed across the top row of the panel (green lines) according to their mean relative frequency at day 0 (from low to high frequency). All of the barcodes identified in resistant CFU are assigned a specific color and position that is aligned in the figure according to their mean frequency in drug susceptible day 0 wells. Both DP and RP cultures from specific time points are marked by different color background. Barcodes found in resistant DP or FP cultures are indicated. Color and position denote a unique barcode. When resistance barcodes were detected in independent cultures they are shown in the same color and X axis position. The Y axis denotes cultures named with the day of treatment and replicate number. **H**, Venn diagram showing the resistance-associated barcodes detected in Experiment 2 and Experiment 3.

### Drug tolerance preferentially develops in clonal subpopulations marked by resistance-associated barcodes

We used the barcode approach to study the overall clonal trajectory of subpopulations that became drug resistant in at least one study well. We identified all of the barcodes present in CFU cultured on rifampicin containing plates of any DP or RP culture in Experiment 2, naming this type of barcode “resistance-associated”. This revealed 28 resistance-associated barcodes in DP and 93 resistance-associated barcodes in RP cultures (of which 13 barcodes were shared by both conditions). We then determined the distribution and number of resistance-associated barcodes in every DP and RP well in Experiment 2, analyzing the wells which contained any rifampicin resistant CFU separately. First, we examined how the number of CFU represented by each resistance-associated barcode changed over the course of Experiment 2, determining that CFU marked by these barcodes exhibited the same two-phase time-kill kinetics (with similar decreases in the read counts of each barcode over time) as the general bacterial population (Fig. 3C). In contrast to the similar *killing kinetics* of each resistance-associated barcode compared to all other barcodes, we found striking differences in the size of the bottlenecks governing the *complete disappearance* of each resistance-associated barcode compared to all other barcodes over the course of the experiment. More specifically, the numbers of different DP resistance-associated barcodes detected at each time point fell at a much slower rate throughout the experiment, decreasing by only 43% over 20 days (Fig. 3D) compared to the numbers of different non-resistance-associated barcodes, which fell by 98% during drug treatment (difference in barcode elimination *P* < 0.0001, estimated by simple linear regression for the slopes). We repeated this analysis for RP resistance-associated barcodes in DD cells and found similar results (*P* = 0.0168) (Fig. 3E). As might be predicted from these results, we also found that resistance-associated barcodes identified during DP were strongly associated with drug tolerance. For Experiment 2, 13 (3%) of the 427 barcodes remaining in day 16-20 drug persistence phase wells were resistance-associated, even after eliminating wells that contained drug resistant CFU (Fig. 3F). In contrast, resistance-associated barcodes only comprised 9 (0.0056%) of the 1,615 barcodes detected in the day 0 cultures (*P* = 0.001, Fisher’s exact test). In summary, we observed that resistance associated barcodes exhibit normal killing kinetics but have a much larger bottleneck governing the *disappearance* of each barcode over the course of each experiment. Together these observations strongly suggest that resistance-associated barcodes do not start out as more tolerant than the majority population, but they appear to acquire drug tolerance more consistently, since they are rarely eradicated over time, even in wells that do not contain any rifampicin resistance.

### Rifampicin resistance emerges repeatedly from an identifiable subset of drug tolerant cells

We found that rifampicin resistance developed repeatedly from the same clonal subpopulations as marked by resistance-associated barcodes. Of the 1,615 barcodes that were inoculated into day 0 wells of Experiment 2, only 28 (1.73%) were resistance-associated on DP. However, 11/28 (39.3%) of the resistance-associated barcodes independently developed rifampicin resistance as shown by their detection in at least two separate cultures of rifampicin-resistant CFU (Fig. 3G). Four of these barcodes were detected in ≥50% of all DP resistant wells and two of these were present in all of the wells that contained drug resistant CFU (Fig. 3G). Similar observations were made in Experiment 3 (Supplementary Fig. 6). It is important to note that these data points represent cultures that were completely independent of each other once they had been seeded into wells at day 0, and that this analysis separately analyzed all wells that ultimately produced rifampicin resistant CFU (Fig. 3A, red dots). Thus, the distinct stability of subpopulations marked by resistance-associated barcodes is unlikely to be due to drug resistance *per se* but is more likely to be a marker for a particular type of drug tolerance that has a propensity for developing drug resistance. Given this likelihood, when did this tolerance and resistance-associated propensity arise? We compared the resistance-associated barcodes from Experiment 2 with those identified in Experiment 3. The two sets of barcodes were completely different (Fig. 3H), indicating that the cells had become predisposed to acquire rifampicin resistance after they had been aliquoted into seed cultures and frozen down for later use, but before they were distributed from these thawed and regrown seed cultures into the wells of the TTR system and exposed to rifampicin.

### Unfixed resistance mutants emerge during the persistence phase

Approximately 95% of clinical rifampicin resistant *M. tuberculosis* strains harbor mutations within the 81 base pair rifampicin resistance determining region (RRDR) of the *rpoB* gene (22). We traced the emergence of each RRDR mutant in Experiment 2 by deep sequencing this locus in colonies isolated from all drug-free and rifampicin agar plates using an experimentally derived cut-off for distinguishing likely resistance mutations from both sequencing and rifampicin induced mutations outside of the *rpoB* gene (Supplementary Fig. 7). Few mutations occurred during early time points followed by an increasing cloud of unfixed mutations in each culture well (Fig. 4A, Supplementary Fig. 8). This heterogeneity in RRDR read sequences predominated by day 20, where 14 out of 16 wells sequenced from drug free plates revealed multiple RRDR mutant alleles above the threshold, even though the WT allele still prevailed in 12 out of 16 wells and only 4 out of 16 wells showed resistant colonies on rifampicin plates. RRDR deep sequencing of the colonies isolated from wells directly plated on rifampicin revealed a single majority resistance allele, but also the continued presence of multiple minority RRDR mutant alleles in cultures from every well that produced resistant colonies (Fig. 4B). A similar picture was seen in studies of plated regrowth cultures, except that significant numbers of mutant RRDR reads appeared at earlier time points (Fig. 4C, D). Of the 35 non-synonymous RRDR mutations detected in this experiment, 19 (54%) have been previously reported in association with rifampicin resistance in clinical or laboratory *M. tuberculosis* isolates (23) (Supplementary Table 1), three were detected only in association with other primary mutations and the remaining 13 were found at low frequency. These results indicate the independent emergence of diverse unfixed RRDR mutations during the drug persistence phase followed by fixation of a single majority resistance allele.

**Fig. 4.**
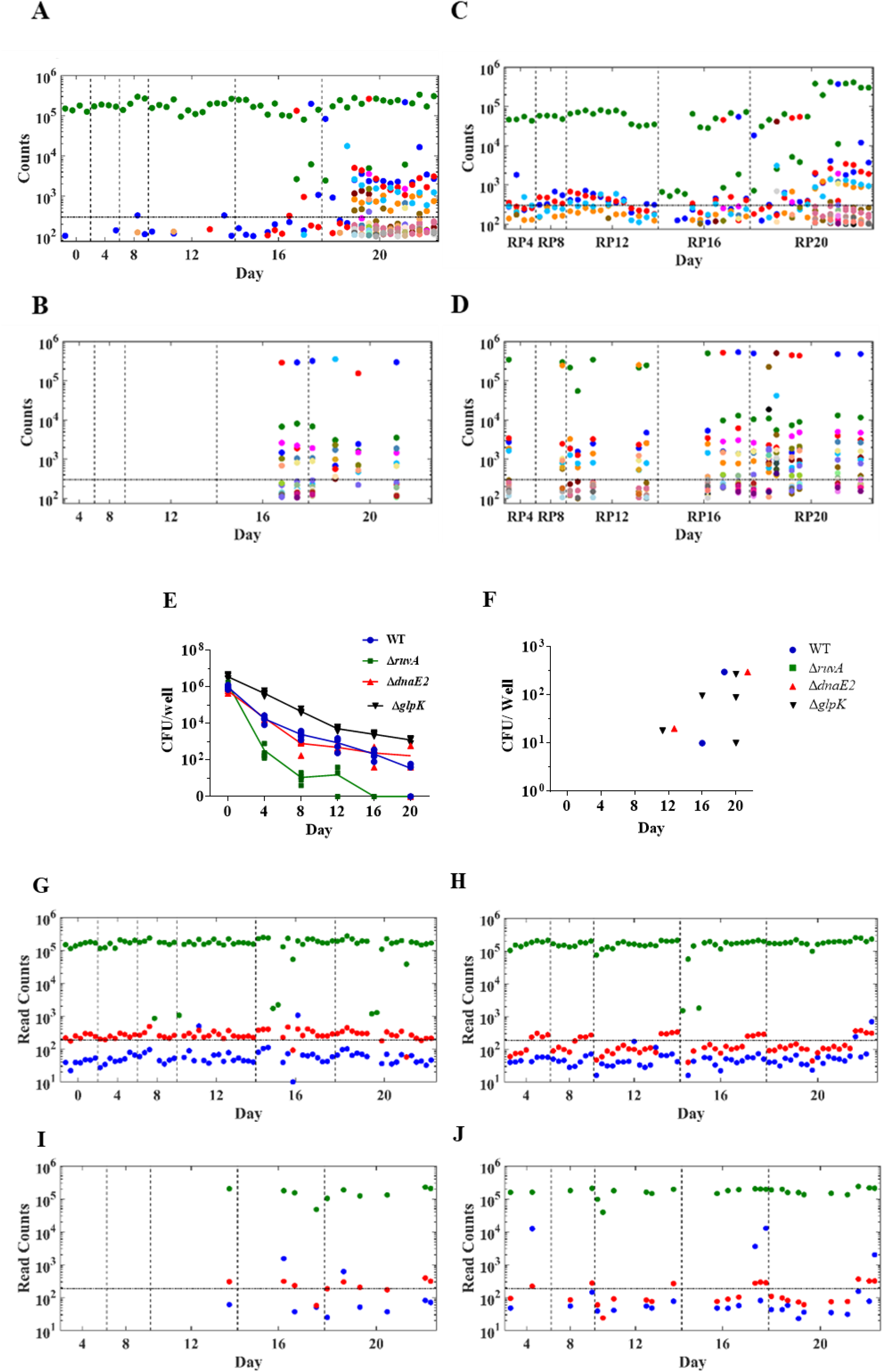
Multiclonal emergence of rifampicin resistance and genetic correlates. **A**, **B**, Rifampin resistance determining region (RRDR) mutations from replicate cultures of direct plated (DP) wells on **A**, drug-free and **B**, rifampicin agar medium and then deep sequenced after amplification with RRDR specific primers. The Y axes show the read count of each mutation detected at different time points. Cultures on drug-free medium from early time points exclusively contained wild type (WT) sequence (green dots) while later time points were predominated by a diverse set of RRDR variants (the full color code key for each mutation is shown in Supplementary Fig. 9). Repeating this analysis on (regrowth plated) RP cultures plated on **C**, drug free and **D**, rifampicin agar medium showed additional RRDR mutants. Dotted line on Y axis denotes the high stringency cutoff for valid read counts. **E**, **F**, Analysis of wild type *M. tuberculosis* and *ΔruvA, ΔdnaE2* and *ΔglpK* knock-out mutants using the TTR method. The CFU/well of each mutant compared to WT control plated on **E**, drug free and **F**, rifampicin agar media. **G-J**, Deep sequence analysis of the *M. tuberculosis glpK* homopolymeric tract (HT) of each culture well and time point in both DP and RP cultures plated on drug free and rifampicin containing media using the results of both Experiment 2 and Experiment 3. The read counts of 6C, 7C and 8C HT variants are represented by red, green and blue dots, respectively. Read counts are shown after DP on **G**, drug free and **I**, rifampicin agar medium and after RP on **H**, drug free and **J**, rifampicin agar medium. The dotted line on Y axis denotes the high stringency cutoff for valid read counts.

### Genetic confirmation that drug tolerance is an important predecessor to drug resistance

We questioned whether drug tolerance must precede drug resistance. The bacterial *ruvA* Holliday junction resolvase gene (24) is one of the SOS response genes that has been strongly implicated as necessary for tolerance to multiple drugs in rapidly growing bacteria (25, 26), and a full deletion mutant of the *M. tuberculosis ruvA* gene Rv2593c (strain *210::ΔruvA)* showed similar MICs towards rifampicin, slightly slower growth in the absence of drug-stress but hypersusceptibility to UV light and DNA damaging agents, without a change in the frequency of rifampicin resistance mutants (Supplementary Fig. 9A-D). We tested a similar *ΔruvA* mutant (H37RvMc^2^6230::*ΔruvA*) in the TTR system (Experiment 4). This *ΔruvA* mutant differed remarkably from WT cells in its ability to survive exposure to 10X the rifampicin MIC (Fig. 4E) and to develop drug resistance (Fig. 4F). All culture wells containing this mutant were sterilized and no drug resistant mutants were detected. An alternative explanation for the development of drug resistance has been the induction of the error prone *M. tuberculosis* DNA polymerase dnaE2(27). In contrast to the *ΔruvA* mutant, similar studies of a complete *dnaE2* deletion (H37RvMc^2^6230::*ΔdnαE2*) showed a two phase kill curve, a drug tolerant phase, and the emergence of rifampicin resistant mutants identical to the parental H37Rv strain.

### Drug tolerance and drug resistance are associated with *M. tuberculosis glpK* gene phase variants

Phase variation of the *M. tuberculosis glpK* gene appears to control the expression of multi-drug tolerance in the laboratory and clinical *M. tuberculosis* strains (13, 14). This form of phase variation is caused by rapidly reversible insertion and deletion (in-dels) events at a 7-C homopolymeric track (HT) of *M. tuberculosis glpK,* which results in reversible frame-shit inactivation of the GlpK protein and activation of a transcriptional program typical for drug tolerant cells (14). We deep sequenced the *glpK* HT in DP and RP cultures on drug-free and rifampicin containing media. The majority of cultures contained bacteria with a predominantly WT 7-C HT along with a small fraction of HT mutants with a single C deletion (from 7-C to 6-C) that was above the mutation threshold (Supplementary Fig. 7A-D). A 6-C *glpK* HT genotype has not been observed in clinical strains. However, we noted that the clinically relevant 8-C *glpK* HT frameshift genotype emerged above background with increased frequency starting on day 12 during the tolerant phase of the killing curve of both the DP and RP wells, where five (two in the DP and three in the RP wells) out of the 104 total wells (4.8%) of the day 12 – day 20 tolerant phase wells showed 8-C *glpK* HT mutants above the limit of detection compared to zero 8-C *glpK* HT mutants emerging in the 32 DP and RP wells plated during days 4 – 8 (Fig. 4G and H), although this difference did not reach statistical significance (*P* = 0.4). In contrast, the 8-C *glpK* HT mutants were significantly enriched in the rifampicin resistant colonies recovered when the DP and RP wells were plated on rifampicin media, where six of the 36 (17%) DP and RP wells that produced rifampicin resistant colonies contained 8-C *glpk* HT mutants above background (Fig. 4I and J) compared to the 4% frequency of 8-C *glpK* HT mutants in the DP and PR wells plated on drug free media (*P* = 0.019, Fisher’s exact test). *M. tuberculosis* with an 8-C *glpK* HT track sequence grows slowly as small colony variants and rapidly reverts to in-frame 7-C *glpK* HT track large colonies under growth permissive conditions, such as are likely to exist once bacteria develop rifampicin resistance at the RRDR. Thus, the appearance of even a minority of 8-C HT reads indicates a strong selection for this genotype in the drug persistent *M. tuberculosis* cells evolving to drug resistance. Consistent with these findings, a complete *glpK* deletion mutant (H37RvMc^2^6230::ΔglpK) demonstrated significantly higher levels of drug tolerance compared to the WT (*P* = 0.002), *ΔruvA* (*P* = 0.0006) and *ΔdnaE2* (*P* = 0.001) mutants (two-tailed unpaired t test) (Fig. 4E). We also compared rifampicin killing at day 4, when all strains had at least some measurable CFUs, and again the *ΔglpK* mutants showed significantly greater survival (Fig. 4F). The emergence of rifampicin resistant mutants during the TTR assay was also significantly increased in the *ΔglpK* mutant compared to the *ΔruvA* mutants (P = 0.037, Fisher’s exact test). There was also more rifampicin resistance detected in the *ΔglpK* mutant compared to the WT, and *ΔdnaE2* mutants; however, these differences were not statistically significant (*P* = 0.42 for both WT and *ΔdnaE2* mutant).

## Discussion

Our results show that certain barcodes are repeatedly selected from multiple cultures by rifampicin treatment. This strongly suggests that the tolerant phenotype can be inherited. However, the observation that a distinctive set of tolerant barcodes arise from each frozen aliquot of the same stock culture emphasizes that this phenotype is also transient and condition dependent. Drug tolerance appears to develop in at least two phenotypically different (DP and DD) subpopulations and resistance-associated barcodes appear to identify subpopulations that acquire drug tolerance at particularly high frequencies. Bacteria with resistance-associated barcodes are able to avoid complete elimination by drug treatment and they then go on to develop drug resistance. Together, our findings suggest that rifampicin resistant *M. tuberculosis* starts out in subpopulations with a special ability to generate drug tolerant clones.

In addition to DP, DD and resistance associated barcoded subpopulations, our results also suggest that another type of drug tolerant subpopulation might exist which is DD, but only by direct plating. The presence of DD cells that can be detected by direct plating but not during incubation in drug free media is supported by our observation that fewer barcodes were recovered in RP cultures than in DP cultures on days 16 and 20 of our experiments. This was a surprising finding because barcodes recovered by regrowth plating (RP) would be expected to include both barcodes recovered by DP plus barcodes associated with additional DD clones. If correct, this expectation would lead to a larger number of barcodes being recovered in RP cultures than DP cultures, and this was indeed the case through day 12 of our experiments. However, after day 12 we recovered fewer barcodes in RP cultures than in DP cultures. We do not have a definitive explanation for this observation; however, it is possible that some drug tolerant clones, such as those that persist through 16 – 20 days of rifampicin treatment, do not tolerate a rapid transition into drug-free liquid media. If true, this hypothesis would describe a new kind of DD cell, further expanding the different types of potential drug tolerant subpopulations in a drug treated culture.

Drug tolerant subpopulations appear to undergo a complex mutational process during the drug persistence phase which sometimes results in acquired drug resistance. These cells first develop numerous unfixed mutations in the rifampicin resistance encoding region, and resistance may then occur through fixation of a predominant resistance genotype. The unfixed mutations that we observed were not deep sequencing artifacts as we defined cutoffs of minimal read counts needed to identify significant mutations based on the false mutation discovery rate of regions outside of the *rpoB* RRDR region on similar-sized PCR amplicons in wells cultured without and with rifampicin. Our discovery is consistent with prior studies of drug resistance evolution in more rapidly growing bacteria during antibiotic exposure and upon recovery (28–32). However, this type of resistance evolution has previously been shown in bacteria exposed to sub-inhibitory concentrations of antibiotics (33), while our *M. tuberculosis* cultures were incubated at constant concentrations of rifampicin at 10X the MIC, a finding that has serious implications in the treatment of human tuberculosis.

The events that cause subpopulations of *M. tuberculosis* to become predisposed to drug tolerance are not clear. However, we showed that *ruvA,* a key gene involved in bacterial DNA repair and oxidative stress responses, is required for drug treated cultures to survive into the drug persistence phase, further implicating antibiotic induced oxidative stress in *M. tuberculosis* cell death and resistance to this stress as a key feature of drug tolerance and subsequent drug resistance. We also discovered that a fraction of emergent drug resistant cells were *glpK* phase variants. Phase variation leading to reversible inactivation of the *M. tuberculosis glpK* gene has been shown to turn on drug-tolerance associated transcriptional programs (13, 14) and provide resistance to oxidative stress caused by hydrogen peroxide. The reversibility of these mutants makes them difficult to detect by measuring CFU from drug - treated wells, but the appearance of *glpK* mutants, once they are fixed by drug resistance, strongly suggests that *glpK* phase variation is at least one of the underlying mechanisms of drug tolerance in our study.

We demonstrate that small subpopulations within each culture are repeatedly able to survive through the bottleneck imposed by rifampicin treatment – identifying these barcode associated clones as having a heritable component that either leads to drug tolerance before drug exposure or causes a predisposition to become repeatedly drug tolerant. One of our interesting findings was that repeating our study by performing Experiment 3 using a new frozen stock of the same barcoded library led to a completely different subset of barcode clones surviving into the persistence phase. These results strongly suggest that a different subpopulation marked by a different set of barcodes develops in each log phase culture that seeds all of the wells in each experiment. Together these observations delineate both the transient development of the tolerant phenotype, which is likely to be stochastically generated in an identifiable subpopulation each time our bacterial stock culture populations were allowed to enter log phase growth before drug exposure, and a heritable component of drug tolerance observed after this subpopulation was aliquoted into different culture wells and exposed to rifampicin. Our observations do not rule out the possibility that *M. tuberculosis* can also stochastically become drug tolerant during drug exposure. Indeed, this may be the case for barcodes that survive through days 16 and 20 in isolated or small numbers culture wells, suggesting additional pathways to drug tolerance.

Our ability to explore the clonal dynamics of drug tolerant subpopulations was critically dependent on several features of our experimental design including rigorous control of drug concentrations, the combined (Experiments 2 plus 3) study of 75 parallel culture wells which enabled us to examine barcode representation at each time point in cultures that did not of themselves influence the outcome of later time points and starting inoculums well below the expected rifampicin resistance mutation frequency, which prevented confounding due to the emergence of pre-existing resistant mutants. The multi-well format also enabled us to exclude barcodes from wells with emergent resistance when required by our analysis. For biosafety reasons, we chose to perform this study with Mc^2^6230, a defined auxotrophic mutant of H37Rv and it is possible that some of our observations could be confined to this strain. However, Mc26230 has drug MICs including rifampicin that are comparable to H37Rv; thus, we believe that our findings are broadly representative of virulent *M. tuberculosis.* We anticipate that the study of separate subpopulations in *M. tuberculosis* cultures using this approach should aid in the discovery of molecular markers as well as the underlying mechanisms of drug tolerance and emergent drug resistance. Targeting these subpopulations with new therapeutics could shorten TB treatment and prevent resistance emergence, hastening the global eradication of this disease.

## Materials and methods

### Strain and culture methods

All experiments were performed using the defined auxotrophic strain of *M. tuberculosis* H37Rv mc^2^6230 (34), (a kind gift from Prof. William Jacobs Jr., Albert Einstein College of Medicine, Bronx, NY), either in its unmodified version or subjected to additional genetic modifications such as chromosomal barcoding or with gene deletions as indicated. Cells were grown in Middlebrook 7H9 broth containing 0.2% glycerol, 0.05% Tween-80, 10% OADC supplement (BD diagnostics, MD, USA) and 24 μg/ml of calcium pantothenate (Sigma, USA). Primary cultures were grown up to mid-log phase (0.6 OD_595nm_), and then diluted 1:10 in fresh 7H9 and distributed into each transwell (200 μl/well) of a 24 well plate (HTS transwell, Corning) using a Hamilton Star (Nevada, USA) robotic liquid handler in a bio-safety level 2 enclosure. The liquid handler was connected to a LiCONiC (Mauren, Liechtenstein) shaking incubator and a BioTek multi-mode plate reader (Synergy Neo, Vermont, USA) for measuring regrowth in the cultures based on optical density. To expose transwells to rifampicin, 7H9 containing rifampicin (Sigma) at 0.05, 0.1, 0.2 and 0.5 μg/ml corresponding to 5X, 10X, 20X and 50X of the MIC_90_, respectively, was added into each of the basolateral wells (1.2 mL) and incubated at 37 °C at 20 rpm with 90% relative humidity. When a constant level of drug exposure was desired, pre-warmed drug-containing medium was replaced in the basolateral wells regularly for up to 30 days. At every time point, 50% of the culture (100 μl) from each of the time point designated transwells was collected and washed once in fresh 7H9 broth. These cells were then plated in two equal volumes onto 7H11 agar (containing 10% OADC, 0.5% glycerol and 24 μg/ml calcium pantothenate), with and without rifampicin at a final concentration of 1 μg/ml. For regrowth plating, the remaining 50% of the drug exposed culture in transwell was allowed to regrow in the absence of rifampicin by replacing the media from the basolateral well at least four times with prewarmed drug free broth and monitored for regrowth. Once the regrowth of approximately 0.2 OD_595nm_ was observed, equal halves of each culture were plated on 7H11 agar with and without rifampicin as described above.

We performed four sets of experiments using *M. tuberculosis* of Mc^2^6230 strains. Experiment 1 is performed with WT Mc^2^6230 which was exposed to 4 different Rif concentrations of rifampicin at steady state concentrations using TTR system. Experiments 2 and 3 were conducted using different aliquots of barcoded Mc^2^6230 transformant pool. In experiment 2, we used 4 replicates from day 0 till day 8, 12 replicates each for day 12 and day 16 while another 16 replicates for day 20. Experiment 3 contained 3 replicates on day 0 and four replicates each during all other time points of rifampicin exposure. Experiment 4 was performed using knockout mutants for *ruvA, glpK* and *dnaE2* along with WT of Mc^2^6230.

### Assays for membrane permeability

The permeability of the transwell membranes to *M. tuberculosis* was studied by incubating actively growing *M. tuberculosis* cultures in quadruplicate transwells with drug free 7H9 broth in the basolateral well for 24 hours at 37 °C with 20 rpm shaking. To check for any bacterial passage into the basolateral compartment, the entire culture medium from basolateral well was plated onto 7H11 agar. To study drug equilibration time between the transwell and basolateral compartment, 7H9 broth containing methylene blue (0.01%) was added to quadruplicate basolateral wells and the absorbance of the transwell broth was measured for over 6 hours at 37 °C.

### Generating an *M. tuberculosis* barcode library

Barcode oligos were custom synthesized on a high-density microarray slide and eluted to generate a barcoded forward primer pool (MYcroarray, Biodiscovery, LLC, Ann Arbor, MI). Each barcode was designed to differ from all other barcodes by at least two nucleotide bases to exclude any sequencing artifacts. A library of 20,000 oligos were synthesized, each containing a unique 11-base barcode with a 5’ flanking sequence (16-base) complementary to the target plasmid for infusion cloning with the 3’ end compatible for amplification of *hygR* cassette from the pKM342-hyg plasmid (5’ CAATACAACCTATTAATTTCTAGACTCGAGGTACCG 3’). The oligo pool was then used for an asymmetric PCR with limited barcoded primer pool and excess reverse primer using AcuuPrime® PfX polymerase (Invitrogen) to incorporate most of the barcoded oligos in the *hygR* amplicon. For infusion cloning, the integrative plasmid pMV306kan (35), (a kind gift from Prof. William Jacobs Jr.) was double digested with VspI and XhoI (NEB) and the fragment lacking *kanR* was purified from the agarose gel. The barcoded HygR amplicon was then cloned using Infusion cloning kit (Takara Bio) as per the manufacturer’s instructions. The ligated product was then used for transforming the *E. coli* strain provided with the kit and the transformants were selected over LB agar plates with 200 μg/ml of hygromycin. Colony PCR and barcode sequencing of 48 randomly picked transformant colonies showed the presence of unique barcodes in each isolate The colonies (>100,000) from multiple plates were scraped, pooled and used for plasmid isolation using Qiagen midi-prep kit.

Mid-log phase *M. tuberculosis* Mc^2^6230 cultures were used for preparing competent cells by washing three times with 10% glycerol. These cells were then electroporated with 1 μg of the barcoded plasmid pool, in replicates. Cells were recovered in 7H9+OADC+Calcium pantothenate broth for 24 hours at 37 °C and the transformants were selected over hygromycin (Hyg) (50 μg/ml) agar plates. All colonies from multiple plates were collected, pooled and glycerol stocks were made. Although the origional microarry generated approximately 20,000 different barcodes, deep sequencing the barcodes from the plasmid pool isolated from *E. coli* identified a total of 4,401 unique barcodes, which was considered as the starting number of possible barcodes for all experiments.

### Preparing cultures for TTR experiments

For the TTR experiment, glycerol stocks of unbarcoded or barcoded strains were thawed and washed once with fresh 7H9 and inoculated into 7H9+OADC+calcium pantothenate+Hyg and sub-cultured once in fresh media until they reached an 0.6 OD_595nm_. Cultures were then diluted 1:10 in fresh media either with or without rifampicin at the indicated concentrations.

### Deep sequencing and data analysis

For each 7H11 plate with visible growth, all colonies were harvested by scraping the entire plate and then resuspending the scraped material in 1X TE buffer with 4 mg/mL of each Lysozyme and Lipase (Sigma) and then incubated overnight in a 37 °C. Cells were lysed by adding 2% SDS (final concentration) and incubated at 55 °C for 15 minutes followed by DNA isolation by using phenol: chloroform extraction method. The barcode region was amplified from the isolated DNA using forward (5’ TCGTCGGCAGCGTCAGATGTGTATAAGAGACAGCATCATGAACAATAAAACTGTCTGC3’) and reverse (5’GTCTCGTGGGCTCGGAGATGTGTATAAGAGACAGACGATGACGGGCTGGTC3’) primers using Phusion high fidelity polymerase (ThermoFisher Scientific) for 20 cycles, containing template denaturation at 98°C for 15 seconds followed by primer annealing at 68°C for 20 sec and an extension step of 72°C for 6 seconds. The product was gel purified (GeneJET, ThermoFisher Scientific). For rifampicin resistance mutation analysis, the RRDR locus from the same DNA samples was amplified using forward 5’ TCGTCGGCAGCGTCAGATGTGTATAAGAGACAGCGGTG GTCGCCGCGATC 3’ and reverse 5’ GTCTCGTGGGCTCGGAGATGTGTATAAGAGACAGGCACGCTCACTGAC AGACC 3’ primers. The amplicons from each sample were uniquely indexed using Illumina NexteraXT V2 indexing primers by a second PCR of 5-6 cycles. The indexed products were purified using AMPure XP beads (Beckman Coulter), Qubit quantified and pooled at equal concentrations. Paired-end deep sequencing was performed using Illumina MiSeq platform at the Genomic Center, Rutgers New Jersey Medical School.

For deep sequence analysis, the FastQ files were first converted into a string matrix and the 11 base pair barcode sequence was extracted between the two flanking sequences (V1, AAACGTCTTGCTCGAG and V2, GTGGCGGCCGCTCTAGAAC) with 100% sequence identity for both forward and reverse reads using the MATLAB bioinformatics toolbox. Any undefined bases and reads above or below 11 bases were excluded during extraction. The selected barcodes were further filtered through an inventory barcode list and the mismatching sequences were excluded. Selected barcodes with a minimum of 10 read counts were used for further analysis. The read count cutoff was determined based on the reads of samples prepared from a known number of CFUs at varying cutoff limits. To determine the fidelity of deep sequencing, a sequence outside the barcode region was extracted at various lengths and measured for sequence variations. For plotting a clonal kill curve, the frequency of each barcode read was calculated from each sample and multiplied with the observed CFU/well producing an estimated CFU represented by each barcode. Statistical significance was estimated using GraphPad Prism 9. For identifying rifampicin resistant mutations, the RRDR sequence (81 bases) was extracted between the two flanking sequences (V1, CGGTGGTCGCCGCGATCAAGGAGTTCTTC and V2, GGGCCCGGCGGTCTGTCAC GTGAGCGTGC) and all the sequence variants above 100 read counts were considered for further analysis. To analyse the sequence variations on the HT tract of *glpK,* the deep sequence data was processed as mentioned before and the region was extracted between two flanking sequences (V1, TACATCGGTGACATGCAC and V2 CGGCCCGCCGGTCAGATTC) and the read counts of the 6C, 7C and 8C variations was recorded.

### Generation of deletion mutants

The *ruvA, dnaE2* and *glpK* genes were deleted using allelic exchange, as described previously (36). Briefly, 1200 to 1500 base pairs upstream and downstream of *ruvA, dnaE2* and *glpK* were amplified and the PCR products were purified and cloned into p2NIL suicide vector, followed by insertion of PacI cassette containing the *lacZ* and *sacB* genes. All cloning was done in *E. coli* Top 10 (Invitrogen) and the final mutant constructs were confirmed by Sanger sequencing. The recombinant plasmids were used to transform *M. tuberculosis* strains and blue colonies (single crossovers) were isolated and grown on 7H10 agar medium without selection and plated on sucrose–X-Gal plates to select for white mutant colonies (double crossovers). The mutant colonies were screened for deletions in *ruvA*, *dnaE2* or *glpK* genes by PCR.

## Author Contributions

JS, PK and DA designed the experiments. JS performed all TTR experiments. AT performed next-generation sequence data processing and visualization. SL designed the custom barcodes. RS and PK generated and tested the barcode library. HS performed all of the *ruvA* knockout experiments in *M. tuberculosis* strain 210 and provided the constructs for *ruvA* deletion in Mc^2^6230. CL generated *glpK* deletion strain in *M. tuberculosis* Mc^2^6230. SRP assisted in statistical analysis. JS, PK, DA interpreted the results. JS, DA wrote the manuscript.

## Data availability

All deep sequencing raw data files are available on the NCBI Sequence Read Archive. The BioProject numbers for data from Experiment 2 correspond to barcode (PRJNA812682, PRJNA812694), glpK (PRJNA812767, PRJNA812854) and RRDR (PRJNA812868, PRJNA812902) are for samples from DP and RP respectively. Data files from Experiment 3 is available under the BioProject number PRJNA812960.

## Code availability

Code for the deep sequence data analysis can be accessed from https://github.com/jeesms39/JS_DA_TTR_BC_RRDR_GLPK.git.

## Acknowledgements

We thank William R Jacobs Jr. from the Albert Einstein College of Medicine for providing the *M. tuberculosis* strain Mc^2^6230. We also thank Mainul Haque and the Rutgers Genomic Center for performing next generation sequencing of the samples. We thank Véronique Dartois from the Center for Discovery and Innovation, Hackensack Meridian Health, New Jersey, for suggestions on developing the TTR system. We thank Neil Stoker from Centre for Clinical Microbiology, University College London, UK, for critical suggestions and for his aid in editing the manuscript.

